# Application of Good Agricultural Practices (GAP) by the Banana Farmers of Chitwan, Nepal

**DOI:** 10.1101/2020.06.12.148551

**Authors:** Arati Joshi, Dharmendra Kalauni, Ujjal Tiwari

## Abstract

The study was conducted to identify the cultivation practices followed by the banana growers and compares it with the framework of Good Agricultural Practices as recommended by the World Banana Forum. Both qualitative and quantitative approach was used for data collection. The scheduled interview was carried out from 103 banana growers, two focused group discussion was carried out with the banana growers, and two key informant interview was carried out; one with the President of Banana Growers Association, Chitwan and another with Executive Director of Chamber of Commerce and Industry, Chitwan. The findings revealed that GAP related to soil management and fertilization (87.4%) and harvesting and on-farm processing (94.2%) is adopted at a low level. Only 36% of farmers apply an appropriate dose of chemical fertilizer, 28% follow crop rotation and 7.8% perform mulching/ intercropping. About 38% of the respondents do not have any source of irrigation. About 98% of the farmers do not perform any processing, cleaning, storing, and packaging of banana. Further, 55% of farmers do not use appropriate personal protective equipment while applying pesticides. The results showed that there is low awareness regarding GAP among the banana farmers in the Chitwan district. Therefore, the conduction of awareness programs and training related to GAP is recommended. By far, no study has been carried out to analyze the good agricultural practices applied by banana growers in Nepal. This study identified the prevailing gap in farmer’s practices and good agricultural practices.

## 1. INTRODUCTION

After the green revolution, the usage of chemical fertilizer has significantly raised to increase crop production globally. Similarly, with the progress in science and technology, the usage of chemical pesticides for the control of disease, pest, and weed had increased. The wide application of chemical fertilizer and pesticide in the crop is making them unsafe to consume, creating a threat to consumers and the producers. There has been bitter evidence of rejection of Nepalese products from the European Union (THT, 2007). Further, it harms the soil, environment, and impedes the trading of agri-product.

The enforcement of Good Agriculture Practice (GAP) has greater relevance in recent days as it ensures safe crop production, facilitate regional trade through the implementation of common GAP standards in the region and ensure acceptability of fruits and vegetables in the international markets. It helps to produce quality goods with high yield that comply with the standards of national and international regulations. Implementation of GAP ensures environmentally friendly agricultural production, considering human health and welfare, improving the profitability and productivity of the farm. There are several examples of the increase in yield of agricultural produce by the implementation of GAP. Danquah *et al*. (2015) reported that the implementation of GAP increases maize yield by 25-27% and cowpea yield by 8-29% in Ghana. Also, the authors reported that the implementation of GAP increases the benefit-cost ratio of maize and cowpea by 36.1-72.8% and 11.1%-19.5% respectively. Dorji *et al*. (2016) reported that the adoption of GAP by farmers leads to increased yield and income. Bairagi, Mishra, and Giri (2018) reported that a 10-point increase in the adoption of the GAP-index increases farm income by 6.2% and decreases chemical fertilizer usage by 31% for paddy, tomato, lentil and ginger in Nepal.

As a member of the World Trade Organization (WTO), Nepal has adopted the Agreement on the Application of Sanitary and Phyto-sanitary Measures and the Agreement on Technical Barriers to Trade. In response to international food safety and quality concerns, to promote sustainable development and increase the export of agricultural produce, Government of Nepal, Ministry of Agriculture and Livestock Development (MOALD) has prepared NepalGAP (Nepal Good Agricultural Practice) Implementation Directives as the first step towards food safety and trade facilitation (MoAD, 2018). For any farm, that wants to receive recognition or certification of Good Agriculture Practice (GAP) has to apply for it in the respective accreditation institute, which in case of Nepal is Department of Food Technology and Quality Control (DFTQC) and then have to successively have to abide by the rules as mentioned in the NepalGAP Implementation Directive. Upon the successful accomplishment of the standards set by the certification body, the farm receives a certification of GAP (MoAD, 2018).

### 1.1 GAP for banana

Fruits are one of the most vulnerable commodities for transmission of the disease since they are consumed in raw form. Certification of fruit, ensuring its safety for consumption is quite important. In Nepal, the area under fruit has increased by 165% in year 2015/16 compared to the year 2000/01 (Pandey *et al*., 2017). Also, fruits are important for improving the economy of the nation. Among the crops, fruit contributes about 7% to total Agriculture Gross Domestic Product (MoAD, 2016). Banana (*Musa spp.)* is one of the important fruit crops of Nepal which contributes about 24.3% of total fruit production (MoAD, 2017). Banana is such a fruit whose demand is high throughout the year (Paudel & Magar, 2016).

Banana is one of the expanding cash crops in Nepal whose area has been increased by 78.5% in the last six years and production has increased by 103.3% (MoAD, 2017, MoAC, 2011). The major banana growing districts are Morang, Jhapa, Chitwan, and Saptari. Among them, Chitwan is the third largest district in terms of area and fourth-largest in terms of production. In Chitwan, the area under banana cultivation is 1,733 ha (DADO, 2014/15) which has increased by 151.1% in the past six years (MoAD, 2017). Banana farming is flourishing mostly in eastern Chitwan and the living standards of farmers in these areas have also been improved due to this enterprise. Banana farming, which was started on 5.34 ha of land in eastern Chitwan in 2002, has now spread over 1733 ha (ICIMOD, 2015; DADO, 2014/15).

Banana is one of the major fruit crops of Nepal in terms of the potential growing area, production, and domestic consumption. However, the productivity of banana in Nepal is only 16 Mt/ha (MoAD, 2017) that is below the average world productivity i.e. 20 Mt/ha (FAO, 2019). Farmers are facing various hassles during production. The wind is one of the major risk factors of banana growers. Along with it, disease and pest have been causing serious damage to the fruit. Despite the continuous efforts of growers to control the disease and pest, the problem keeps causing nuisance since the planting to the fruiting period. Most of the farmers use sucker or rhizome as the planting materials (CADP, 2008), which passively carry the traces of pathogens from the mother plant. And the poor orchard management practices along with it provide a greater probability of disease and pest occurrence. Greater application of pesticides leads to residue occurrence in the fruit. In Nepal, pesticide residue in fruit crop is found to be 0.029187 kg ai/ha (Adhikari, 2017). The conventionally produced banana gives a low yield and is inferior in quality. Despite the increasing area under banana cultivation, productivity remains the same because of the lack of knowledge about the good agricultural practices among the farmers, and it is making the banana enterprise less profitable. To reduce the existing yield gap, banana farmers must adopt good agricultural practices. Joshi, Kalauni, and Tiwari (2019) reported that only 28.2% of the Banana farmers of Chitwan district were aware of GAP for banana. However, no study has been carried out to analyze the good agricultural practices applied by banana growers. This study will identify the prevailing gap in farmer’s practices and good agricultural practices. Awareness and knowledge of new technologies is the first step toward its adoption (Roger, 1995). Therefore, this study assessed the awareness of GAP among banana farmers.

## 2. METHODOLOGY

The research was conducted in the command area of the PMAMP PIU Banana Zone, Chitwan. The study area was Kalika Municipality (Ward No. 1 to 8), Ratnanagar Municipality (all wards), Khaireni Municipality (all wards) and Bharatpur Metropolitan City (Ward no. 1). A list of banana growing farmers provided by the Banana zone, PMAMP Chitwan was used to randomly select the respondent. The scheduled interview was carried out with 103 banana growers, selecting 10% farmers from each area i.e. 10 from Bharatpur Metropolitan City, 10 from Khaireni Municipality, 36 from Ratnanagar Municipality and 47 from Kalika Municipality. Key informant interviews (KII) and focus group discussion (FGD) were also carried out to triangulate data and information obtained from schedule interview and to obtain additional qualitative information.

The secondary data related to banana production was obtained from different institutes and organizations such as Agribusiness Promotion and Marketing Development Directorate, Ministry of Agriculture and Livestock Development, Central Bureau of Statistics, Agriculture Learning Center, Chitwan, Fruit Development Directorate, Food, and Agriculture Organization, etc. Both the primary and secondary information collected from the field survey and other methods was coded and tabulated on Statistical Package for the Social Sciences (SPSS). The analysis was done through SPSS, Stata and MS-Excel.

The farmer’s perception toward GAP was analyzed using different variables. The perception was analyzed on a scale of strongly agree, agree, neither agree nor disagree, disagree and strongly disagree. Perceptions toward GAP were analyzed using an index of agreement. The frequency of agreement was calculated by the summation of the frequency of response of scale as strongly agree and agree. And the frequency of disagreement was calculated by the summation of the frequency of response of the scale: neither agree nor disagree, disagree and strongly disagree. Index of the agreement was calculated by using the following formula:

Index of agreement = (Frequency of agreement - Frequency of disagreement) / n Where, n = total sample size.

When the value of the index of the agreement is greater than 0.5, the variable is considered to have a positive perception, whereas when the adoption index is less than 0.5, the variable is considered to have a negative perception or attitude.

A GAP framework for banana made by World Banana Forum was used as a standard for evaluating the extent of GAP practices followed by banana growers. Descriptive statistics were used to evaluate the extent of application in percentages. The extent of the GAP application was analyzed under a different category as mentioned by World Banana Forum which is as: Soil Management and Fertilization, Water Stewardship, Crop Production, Crop Protection, Waste, and Energy Management, Harvesting and Farm processing and Workers healthy and safety.

## 3. FINDINGS

### 3.1 Farm Size

The mean banana cultivated land of the study area was 4.1390 (±6.6364) ha, of which average 0.4629 (±0.4484) ha area of own land is used by the banana farmers for cultivating banana. Farmers were found to use 0-2 ha of their land for cultivating banana. Most of the farmers cultivated banana only and only a few of them grow vegetables and cereals. About 71% of farmers took land in the lease for cultivating banana. Average of 3.2231 (±6.7642) ha of land was taken in the lease for cultivating banana. Most of the area about 57.5% of the area does not have the source of irrigation. Average 2.3830 (±5.9447) ha area of banana land was found to be non-irrigated and only 1.7560 (±2.6723) ha area was irrigated. The detail of land use under banana farming is shown in Table 1.

**Table 1.** Land use under banana farming

|  | Own (ha) | Leased in (ha) | Irrigated land (ha) | Non-irrigated land (ha) | Total cultivated (ha) |
| --- | --- | --- | --- | --- | --- |
| Mean | 0.46 | 3.70 | 1.75 | 2.38 | 4.13 |
| Std. Deviation | 0.44 | 6.70 | 2.67 | 5.94 | 6.63 |
| Minimum | 0.00 | 0.00 | 0.00 | 0.00 | 0.10 |
| Maximum | 2.02 | 47.29 | 13.90 | 47.29 | 47.29 |

Source: Field survey, 2019

Shrestha *et al*. (2018) reported that the average landholding of banana farmers of Thimura, Ramnagar, Padampur, Ratnagar and Jagatpur area was 1.96 ha. The authors reported the average banana irrigated area to be 0.22 ha and an average banana unirrigated area to be 0.14 ha.

Similarly, Ghimire, Koirala, Devkota, and Basnet (2019) reported that the average area under banana farming in Ratnanagar and Khaireni municipality was 1.92 ha. The area under banana cultivation in our study is found greater than the past studies. It might be because of the different study areas and greater study area coverage in our case (i.e. Kalika Municipality, Ratnanagar Municipality, Khaireni Municipality, and Bharatpur Metropolitan City).

### 3.2 Propagating material used by banana growers

The greater number of farmers used sucker propagated material (95.1%) than tissue culture (1.9%) and only 2.9% used both of them in their orchard (Figure 1). Because of the easy availability of sucker and cheap price, farmers mostly use it. Similarly, CADP (2008) reported that farmers of Morang and Sunsari also used sucker as propagating material to a higher extent. However, due to an increasing infestation of disease and pests, some farmers used tissue culture plantlets.

**Figure 1.**
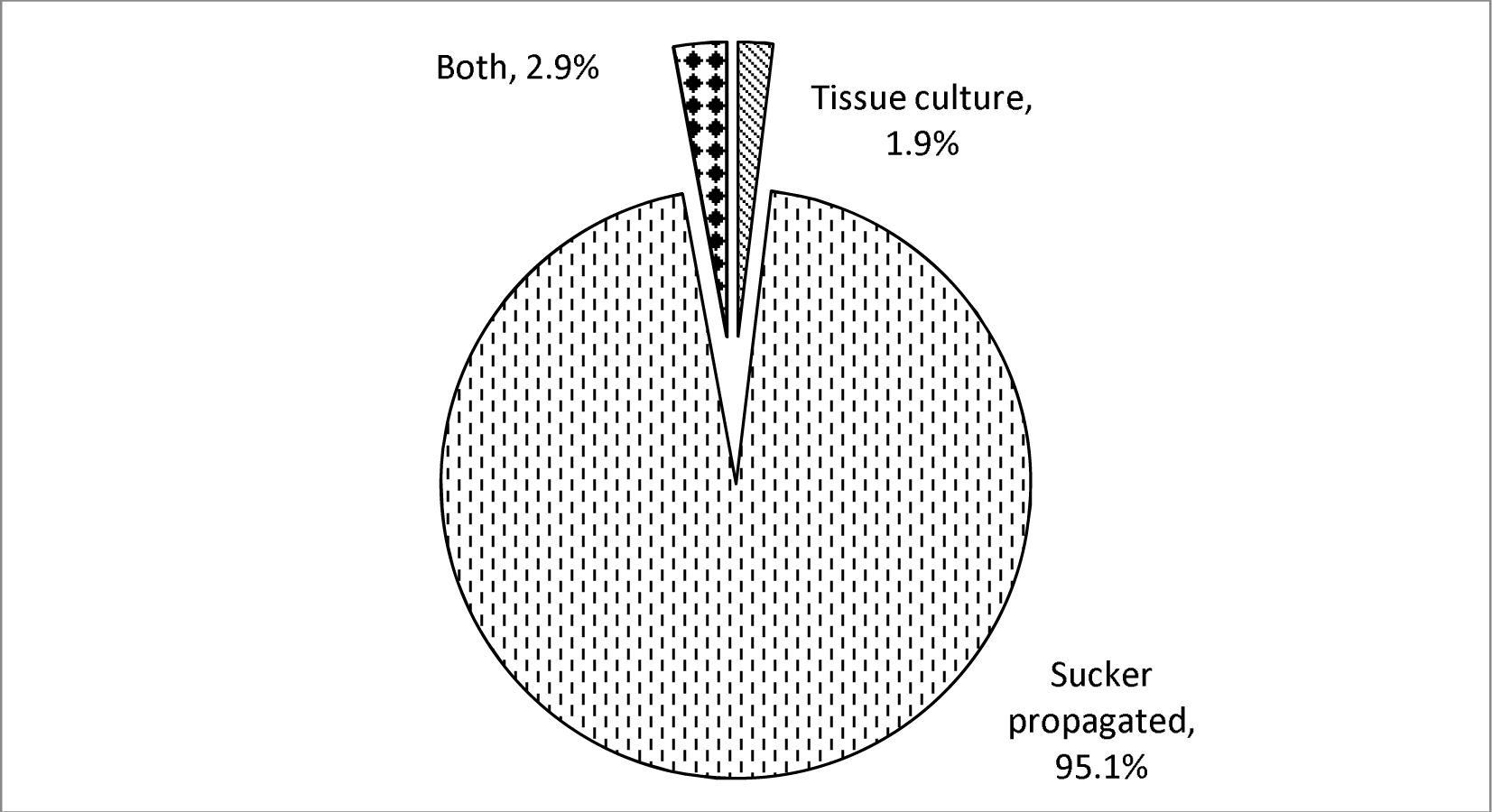
Propagating material used by the respondents

### 3.3 Banana variety used by farmers

Among the total respondents, 95.1% have planted local variety Malbhog, 3.9% planted the variety Grand Nain (G-9) and 1% planted variety William Hybrid (Figure 2). Farmers have received the tissue culture saplings of variety William Hybrid from DADO during the OVOP program. Variety G-9 was imported by farmers from the tissue culture laboratory of Nepaljung and some from Kathmandu. Few farmers explained that the tissue culture plantlet they received was not of good quality. It was a mix of Malbhog and G-9. Farmers have to face a heavy loss due to the poor quality of tissue culture sapling because such a plant got severely infested with disease and pest in the later stage. CADP (2008) reported that progressive farmers prefer tissue culture saplings, but unavailability of it locally in Sunsari is the major problem. A similar problem was observed in the Kailali district (NHPC, 2017). Therefore, most of the farmers have shifted to the use of local variety Malbhog. In our study, the respondent gave a higher preference to local variety, Malbhog for its high price and demand, drought-tolerant ability and high keeping quality.

**Figure 2.**
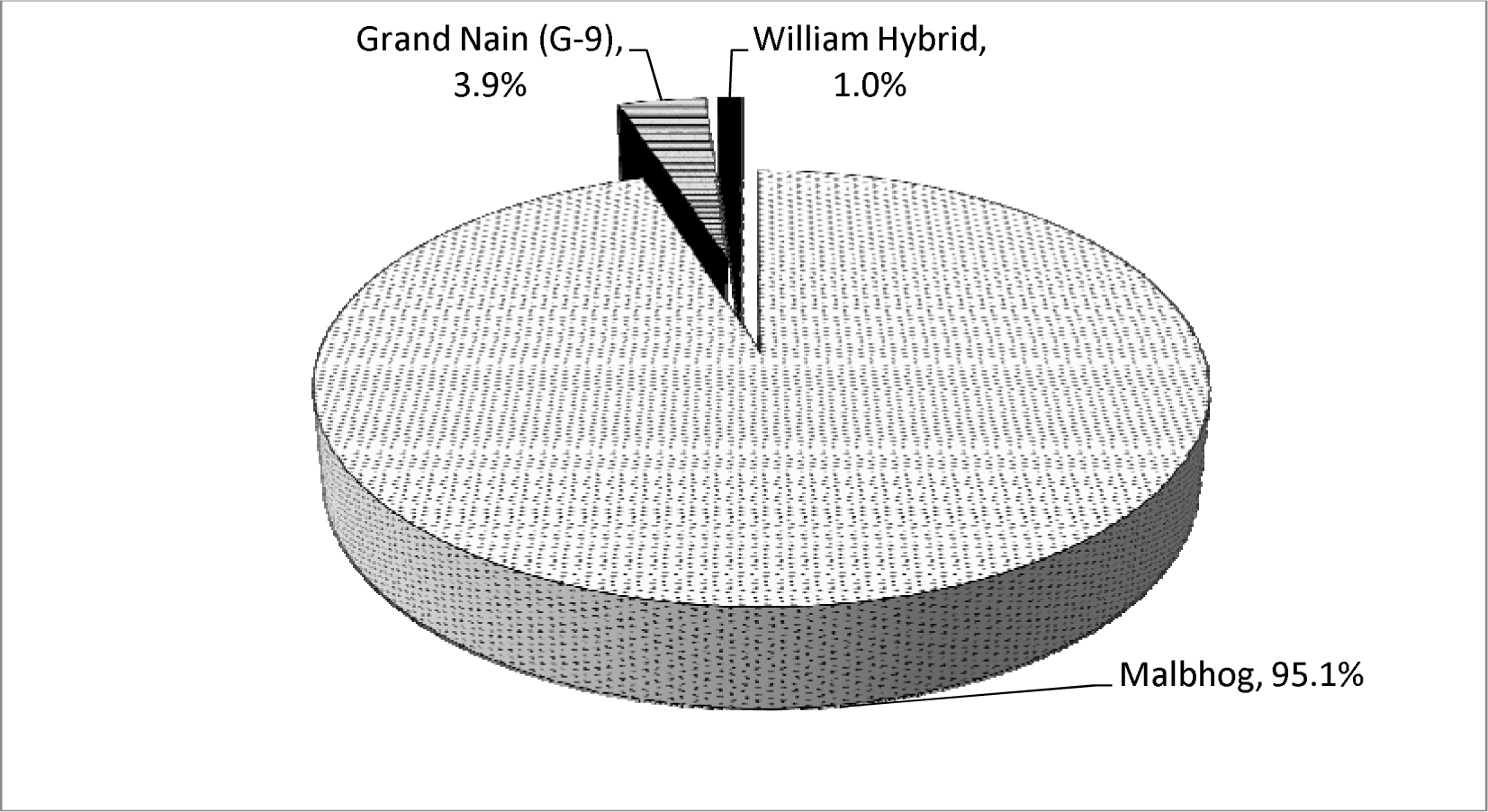
Banana variety used by growers

### 3.4 Perception of farmers toward GAP

The perception of respondents towards GAP is analyzed using an index of agreement. The value of an index of agreement may vary from -1 to 1, with values greater than 0.5 indicating positive responses. The perception of respondents toward GAP was found positive except for the statement; GAP reduces the input cost of production. About 61.2% of the respondent (index= - 0.22) were found to disagree with this statement. While the entire respondent agreed to the fact that, GAP is a sustainable practice, GAP produced crop has higher quality, and GAP reduces farmers’ exposure to health hazards. About 83.3% of the respondents (index = 0.66) agreed to the statement that GAP is a time-consuming practice. Respondent stated that GAP requires high care and time for management. Similarly, 89% of the respondent (index = 0.77) agreed that GAP reduces all forms of pollution. About 83.3% of the respondent (index = 0.67) agreed that GAP increases the farmer’s income. The perception of respondents regarding the other statement is shown in Table 2.

**Table 2.** Perception of respondents toward GAP

| Statement | Strongly Agree (%) | Agree (%) | Neutral (%) | Disagree (%) | Strongly Disagree (%) | Index of agreement |
| --- | --- | --- | --- | --- | --- | --- |
| GAP is a sustainable practice for future | 83.3 | 16.7 | 0.0 | 0.0 | 0.0 | 1 |
| GAP produced banana has nicer appearance and quality | 94.4 | 5.6 | 0.0 | 0.0 | 0.0 | 1 |
| GAP is a time consuming practice | 72.2 | 11.1 | 0.0 | 16.7 | 0.0 | 0.66 |
| GAP reduces all form of pollution | 77.8 | 11.1 | 11.1 | 0 | 0.0 | 0.77 |
| GAP reduces cost of production | 33.3 | 5.6 | 0.0 | 55.6 | 5.6 | -0.22 |
| GAP increases farmer's income | 61.1 | 22.2 | 11.1 | 5.1 | 0.0 | 0.67 |
| GAP reduces farmer's exposure to health hazards | 72.2 | 27.8 | 0.0 | 0.0 | 0.0 | 1 |

Source: Field survey, 2019

### 3.5 GAP application by farmers

#### 3.5.1 Soil management and fertilization

Only 35.9% of the respondents applied the appropriate amount of fertilizer to the banana crop (Figure 3). Most of the farmers applied the organic manure during land preparation, and the chemical fertilizers like urea, potash, and DAP were applied once the plant gets established, i.e. after one month of planting. Only a few farmers top-dress the fertilizers in a recommended interval, while most of them only apply the fertilizer once. Only 32.5%-50% of farmers applied the recommended dose of fertilizer in banana in India (Hassan, 2016). About 28% of the respondent followed crop rotation, while most of the farmers replaced the field with another variety of banana or abandon the field taken on lease. Tiwari *et al*. (2006) reported that only 12% of banana growers of Chitwan follow crop rotation. Only 7.8% of the farmers planted a cover crop or intercrop between the banana inter-rows. Respondents plant vegetables like-cauliflower, bean, and others during the early stage of the plant. Hassan (2016) reported that only 2.5-5% of farmers have practiced intercropping in banana in the early stage in India.

**Figure 3.**
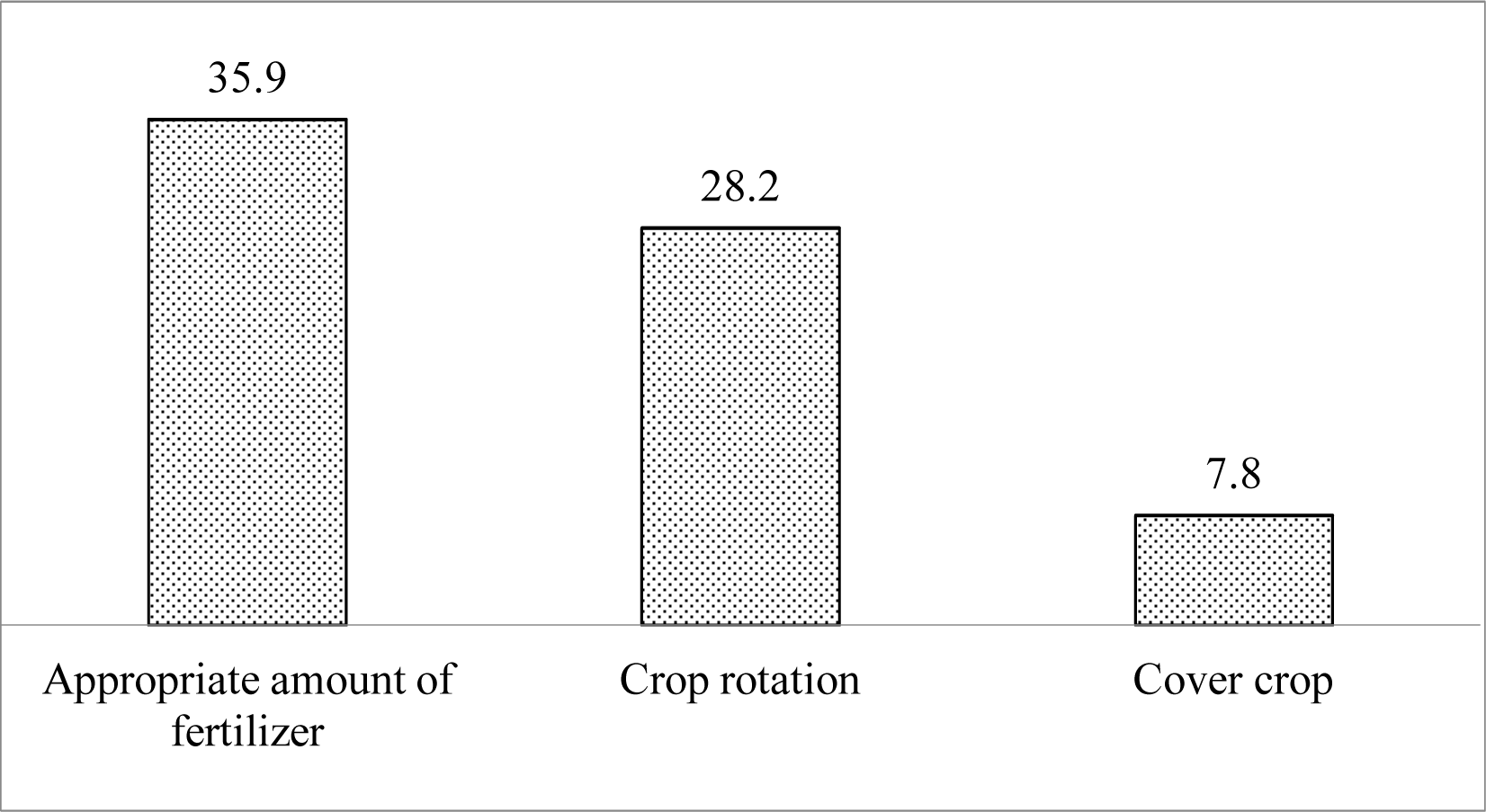
Percentage of respondent farmers adhering to GAP related to soil management and fertilization

#### 3.5.2 Water stewardship

About 91% of the respondents had a drainage facility in the banana orchard and there was no water logging problem (Figure 4). Similarly, 92.2% of the respondents reported that they used organic matter in the soil while planting banana. Organic matter helps to hold moisture in the soil. About 37.9% of the respondents do not have any source of irrigation in their orchard and depend on the rainfall for irrigating their fields. Shrestha and Giri (2012) reported that 37% of farmers of Kailali district faced irrigation problems, and Poudel (2011) reported scarcity of irrigation as the major problem faced by banana growers in Nawalparasi. About 62.1% respondent provided clean and pure water to banana plant that is free from the harmful micro-organism. Most of the farmers irrigate their fields by pumping groundwater and some of them through the canal. About 81.5% of the respondents do not supply irrigation water regularly. Only 2.5% of the farmers have adopted water-saving practices like drip irrigation.

**Figure 4.**
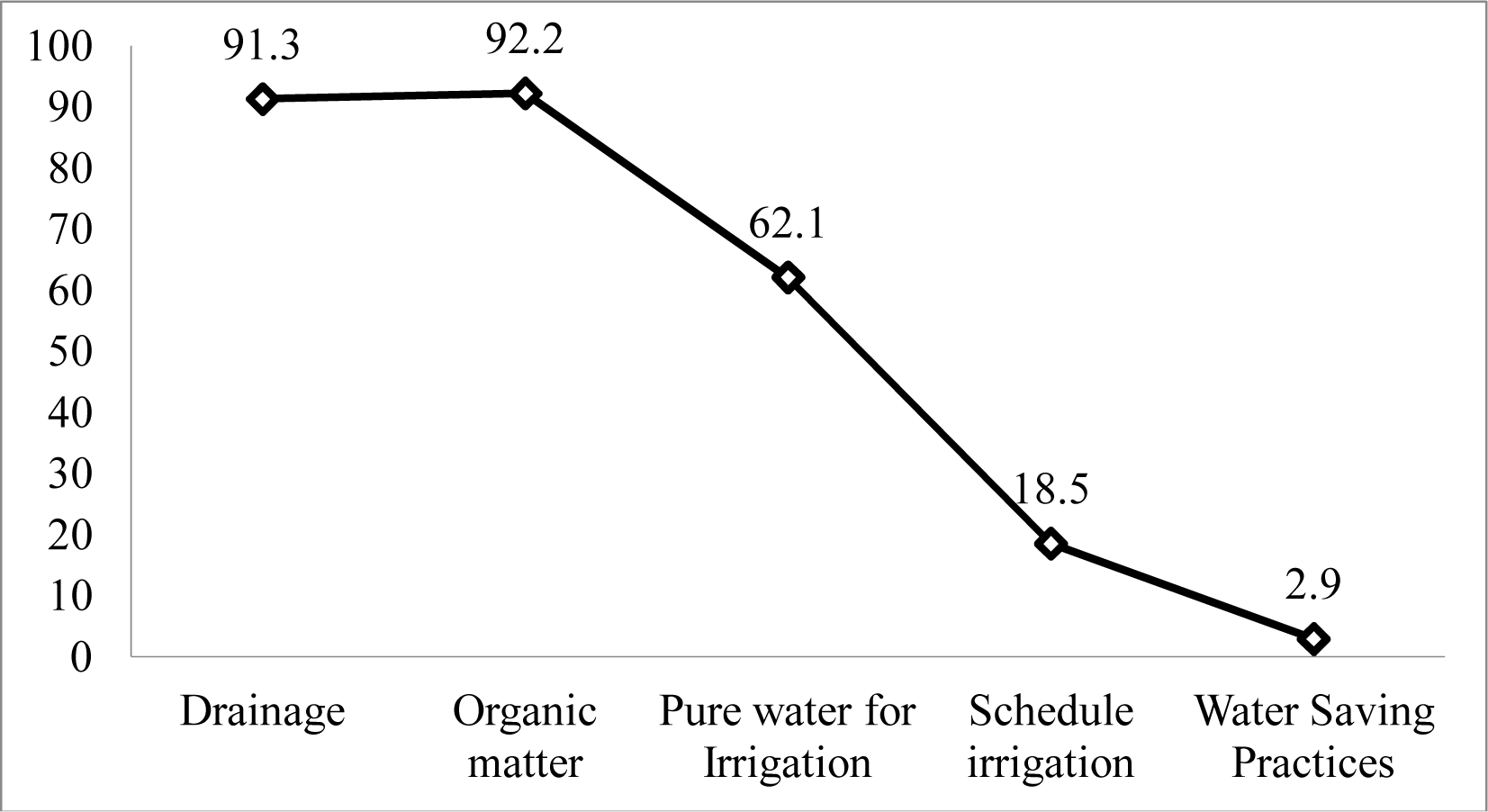
Percentage of farmers adhering to GAP related to water stewardship

#### 3.5.3 Crop production

About 48.5% of the respondents have performed the soil test of their banana orchards (Figure 5). They have performed the soil test during the roving soil test program organized in their area. About 26.2% of the respondents do not apply the fertilizer in a recommended interval. Gautam and Tiwari (2007) reported that applying a higher dose of nitrogen at a time leads to nitrification and leaching, therefore, nitrogen should be applied two months, four months and six months after sucker planting. Similarly, potash should be applied during sucker planting and bunch formation. Only 33% of the potato farmers of Sri Lanka have performed the soil test once in two years and only 26% of farmers apply the recommended fertilizer (Senanayake & Rathnayaka, 2015). Almost all of the farmers except 5.8% were found to recycle the residues (i.e. pseudostem, leaves) of the banana crop as organic manure. About 73.7% of the farmers do not to safely regulate the equipment used in banana farming viz. sickle, shovel and other. Most of them clean the equipment only by water while some of them practice solar drying of the equipment after use, for disinfection.

**Figure 5.**
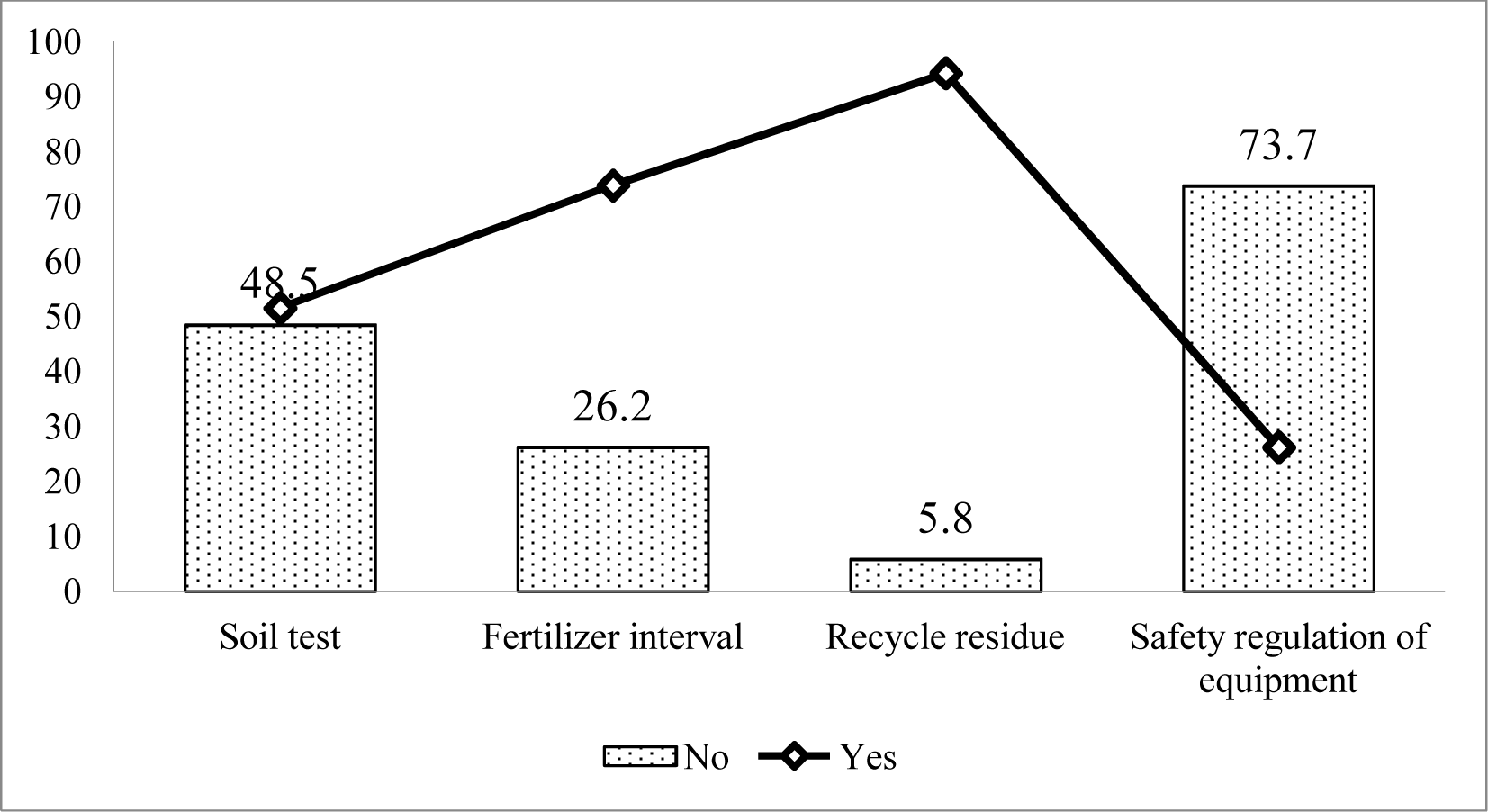
Percentage of farmers adhering to GAP related to crop production

#### 3.5.4 Crop protection

About 74.8% of the farmers applied a chemical pesticide in their orchard based on the weather condition (Figure 6), while others used to mix stickers in the chemical pesticide and apply it in the field irrespective of the weather condition. About 95% of the respondents made a regular survey in their field and implement immediate crop protection measures in case of pest and disease outbreak. However, only 4.5% of the respondents were found to use the disease-resistant variety of banana plant viz. William Hybrid and Grand Nain (G-9). Only 5.8% of the farmers adopted mechanical weed control and only 3.9% of respondents have adopted IPM measure of disease and pest control in their orchard. About 87.4% of the respondents mobilized the experienced worker for handling and application of pesticides. About 35% of the respondents do not know registered or banned pesticides. According to THT (2018), farmers of Dhankuta are using banned pesticides like-monocrotophos, phorate, and quintalphos. Only 44.7% of the respondents used all the personal protective equipment (PPE) while spraying chemical pesticides, while others used only a few of the personal protective equipment (e.g. mask only, mask and boot only). Rijal *et al*. (2018) reported that 34% of farmers use masks only and 14% of farmers do not use PPE at all while spraying the pesticide. About 26.3% of the respondents safely maintain their equipment and 39.8% of respondent keep the record of the use of pesticides.

**Figure 6.**
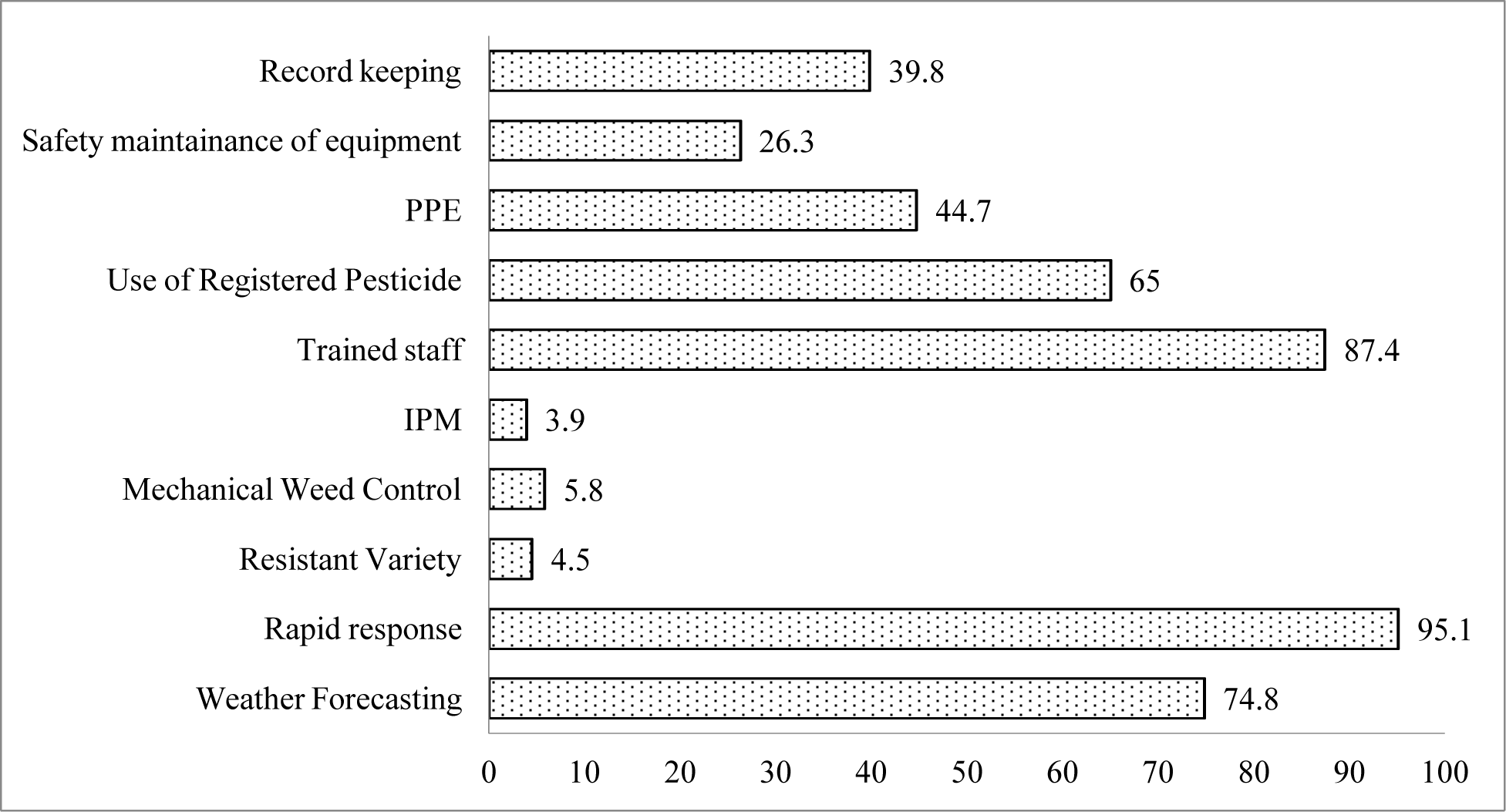
Percentage of farmers adhering to GAP related to crop protection

#### 3.5.5 Harvesting and on-farm processing and storage

About 97.1% of the respondents considered the pre-harvest interval (Figure 7). Usually, farmers do not apply chemical pesticides after finger formation in the bunch. Only 1.9% of the farmers in the study area cleaned the harvested banana and safely stores it in a clean and hygienic condition. Similarly, negligible farmers were found to process the banana and perform packaging before shipment. Farmers of the study area, sell the banana on the orchard itself. Usually, the trader (mostly from Chitwan, Kathmandu, Pokhara, Butwal) come to the farm, harvest the quantity of banana required, load it in the vehicle (usually bolero) and transport it to the destination. NHPC (2017) reported that while transporting banana no packaging materials are used except covering by the banana leaves in Kailaki district. This may cause physical injury to the banana fruits. The post-harvest loss in a banana is about 10-15%, which is greater than mango, citrus, and apple. Noncompliance with the post-harvest practices may lead to the infestation of the pathogenic micro-organism. Khadka *et al*. (2017) reported that the adoption of post-harvest handling practices in tomato, reduces the aerobic bacterial count, coliform bacterial count, mold growth and increases the self-life.

**Figure 7.**
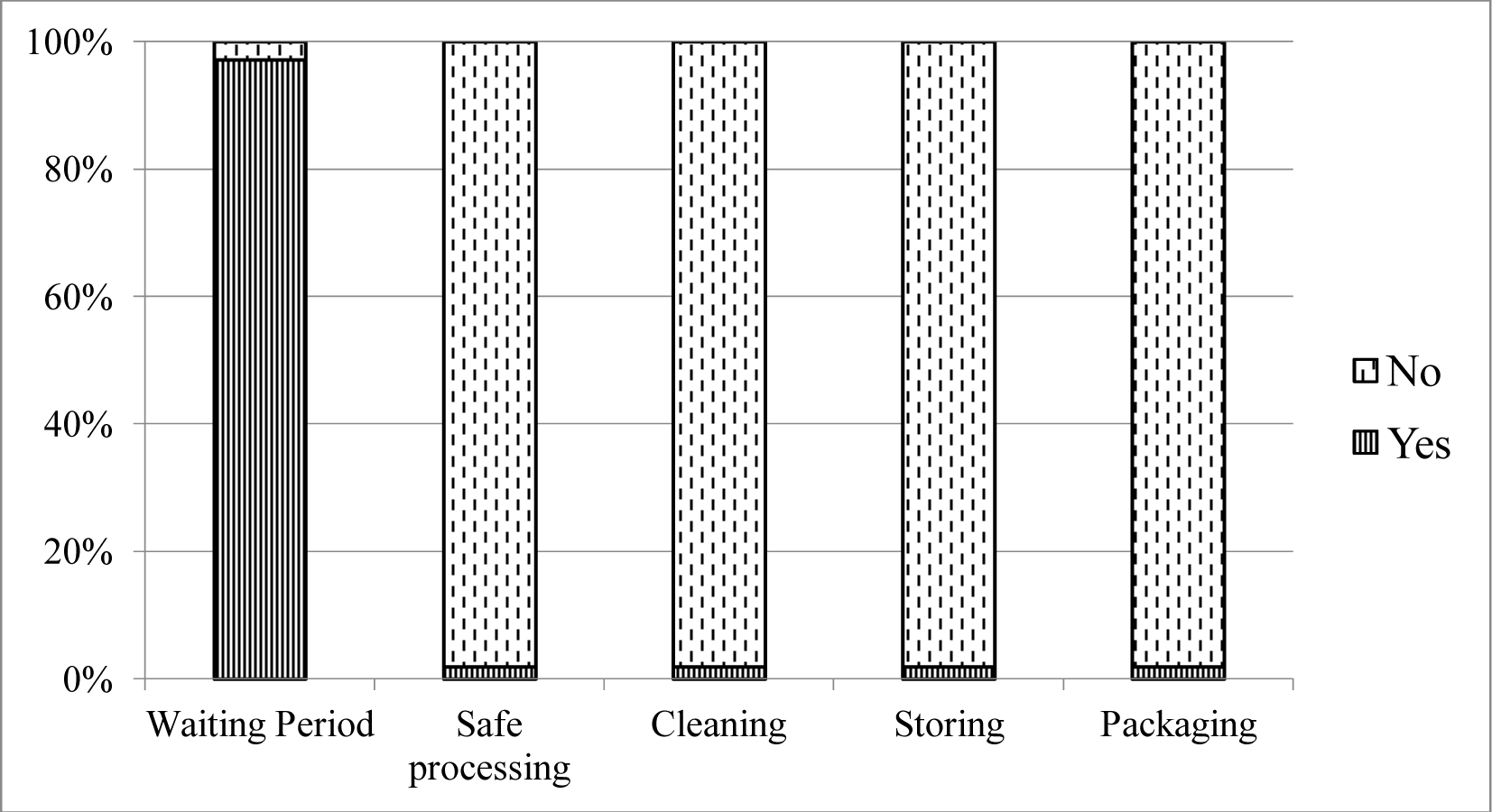
Percentage of farmers adhering to GAP related to harvesting and on farm processing and storage

#### 3.5.6 Energy and waste management

The entire respondent safely disposes of the farm wastes like-nutrients, empty pesticide containers, tanks, and others. Most of them send such waste with the garbage collecting vehicle of Municipality/ Metropolitan City. Some of them bury such waste in the pit while some burn it. The respondent took care of the fact that such waste must not affect the children or animals. About 85.4% of the respondent securely stored fertilizer in a safe place (Figure 8). About 43.2% of the farmers reported that they have been trying to minimize the non-recyclable waste in the farm and recycle the organic and inorganic waste. Only 17.5% of the farmers recorded the total energy consumption on the farm, while none of the farmers have adopted alternative energy sources and establish emergency procedures to limit the risk of pollution. CADP (2008) reported that in Morang and Sunsari, a high share of costs incurred in banana cultivation is from irrigation, which is done by diesel-powered pumping set. The dependency of banana cultivation on such non-renewable fossil fuel does not primarily increase the cost of production but also affects the environment. Some alternatives might be rainwater harvesting, use of solar energy for water pumping, multiple uses of water and drip and sprinkler irrigation technology.

**Figure 8.**
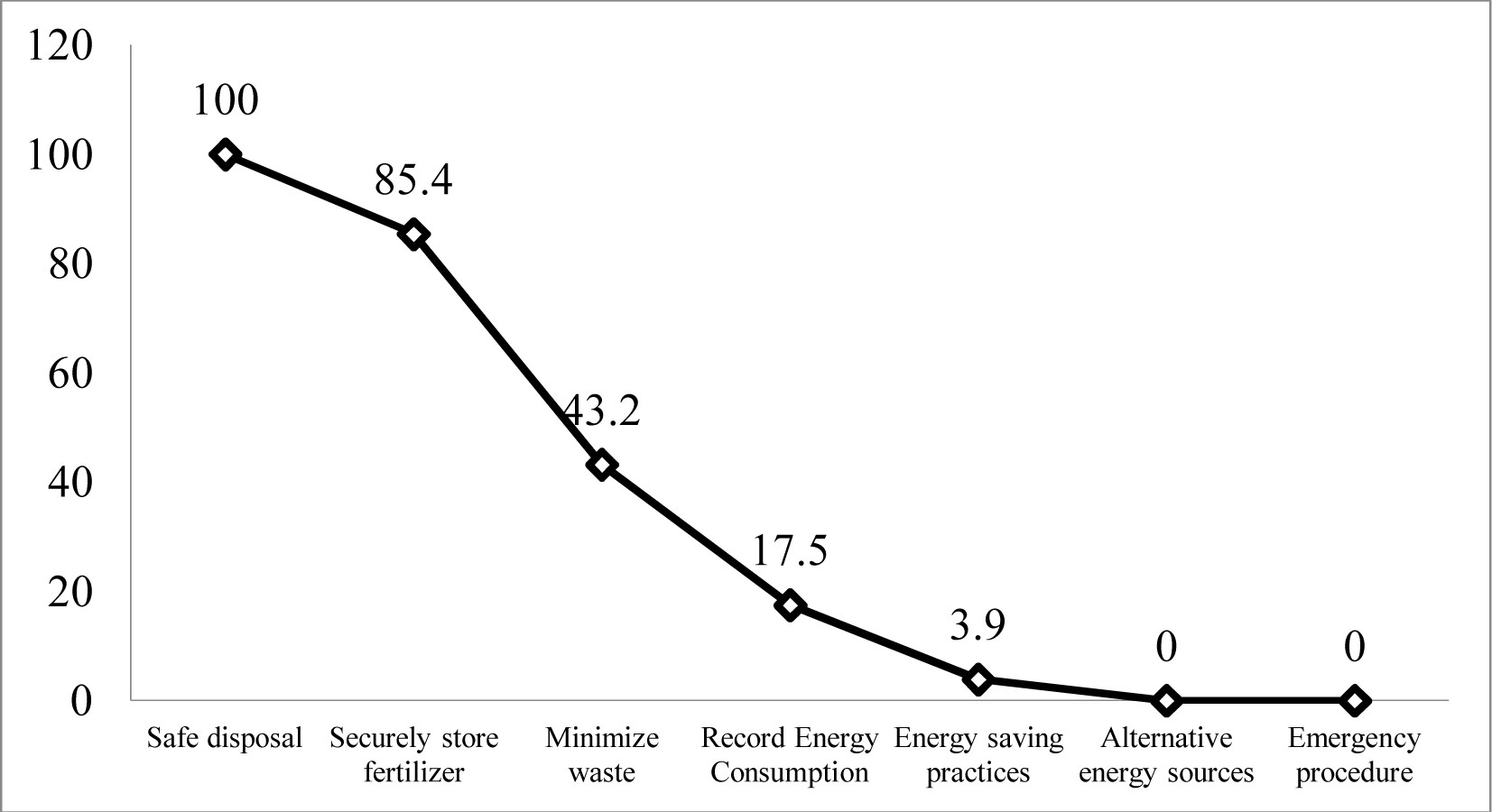
Percentage of farmers adhering to GAP related to energy and waste management

#### 3.5.7 Human welfare and health and safety

About 88% of the respondents have trained the worker to safely use the tools used in the banana farm (Figure 9). Similarly, about 82% of the respondents have provided decent wages to the

**Figure 9.**
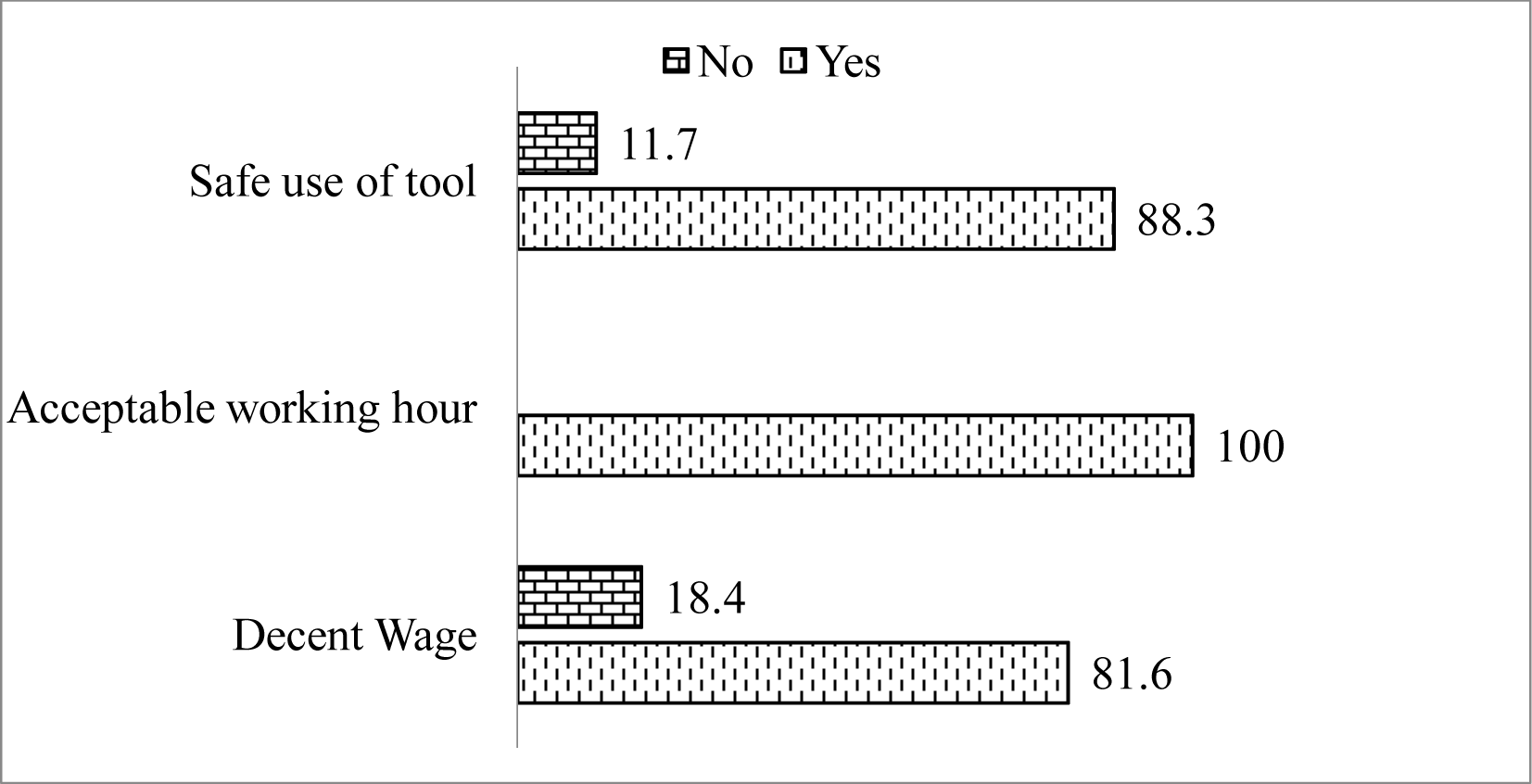
Percentage of fanners adhering to GAP related to human welfare and health and safety

workers based on the intensity of work. The entire respondent reported to provide an acceptable working hour for the workers with the proper rest time.

### 3.6 Total GAP practiced by the respondent

Based on the mean of GAP application (i.e. 0.4), the category of GAP practiced is classified as none applied (0), low level of application (0.1-0.3), medium level of application (0.4-0.7) and high level of application (0.8-1) as shown in Figure 10. About 44% of the good agricultural practices of soil management and fertilization are not practiced by the respondent. Low-level GAP was practiced in harvesting and on-farm processing (94%), energy and waste management (55%), and soil management and fertilization (44%). Medium level of the application was for crop protection (80%) followed by water stewardship (78%) and crop production (74%). The highest level of the application was for elements: human welfare and health and safety (79%), followed by water stewardship (22%) and, crop production (11%).

**Figure 10.**
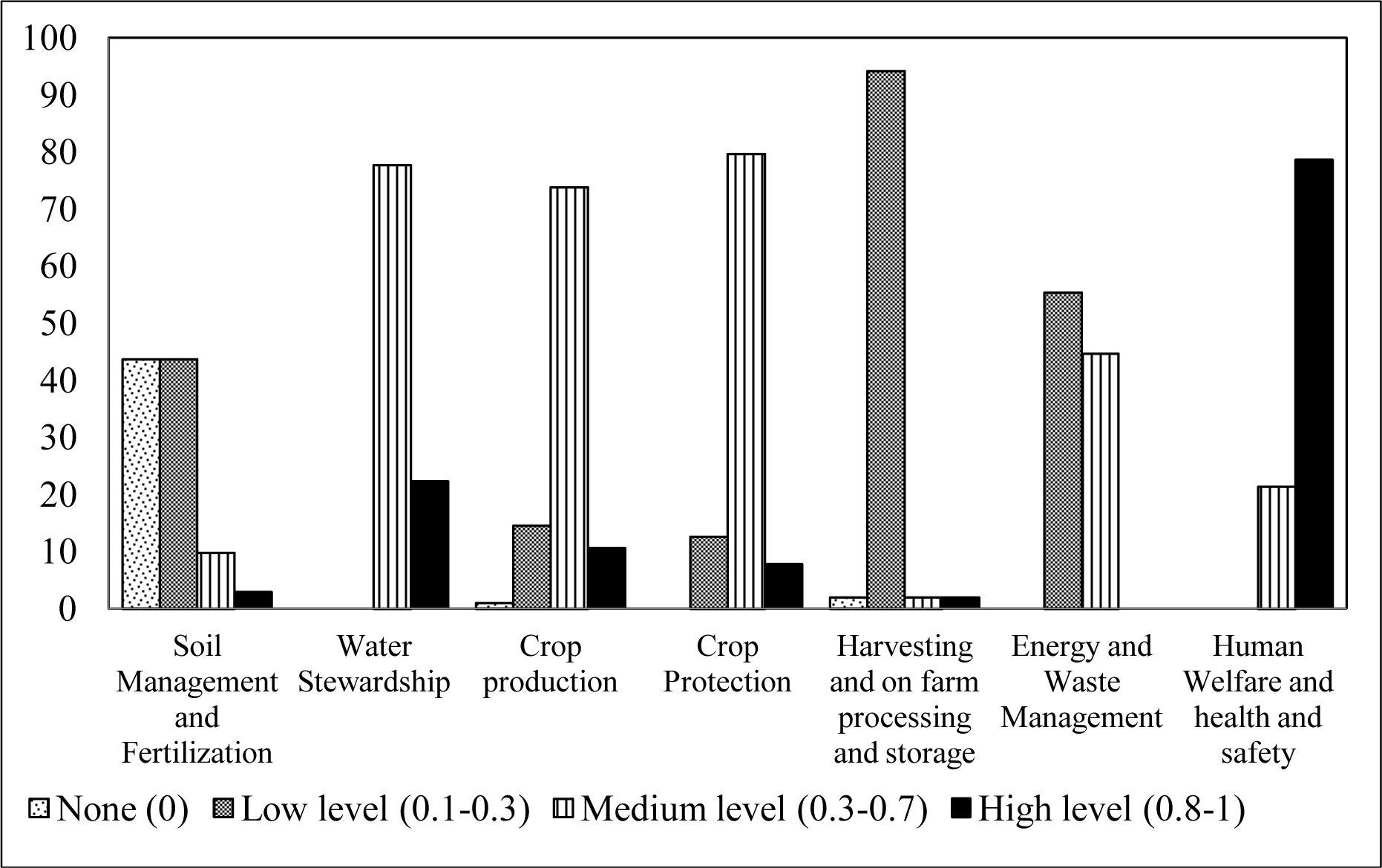
Levels of GAP application by respondent (based on mean of application)

## 4. CONCLUSION

The majority of farmers in the Chitwan district take land in the lease for banana farming. However, most of the banana cultivated area was non-irrigated and therefore, farmers mostly prefer local drought-tolerant variety Malbhog. Farmers of the study area practice poor soil management, harvesting, and post-harvest handling and energy management. To improve the yield and quality of banana in the future, attention should be given to these practices. Good agricultural practice as reported by different researches is a one-stop solution to enhance the production, productivity, and quality of the Banana. Thus to revitalize the Banana subsector, we need to educate farmers about good agricultural practices.

